# Inhibition of Mitochondrial Respiration Fragments ER Architecture and Remodels Organelle Contact Sites, as Revealed by FIB-SEM

**DOI:** 10.64898/2026.05.25.727587

**Authors:** Andrea Dlasková, Bazila Bazila, Pavel Křepelka, Rhea Christine Victor, Darsh Jaresh Jhala, Petr Ježek

## Abstract

The endoplasmic reticulum (ER) and mitochondria maintain a dynamic structural partnership essential for pancreatic β-cell homeostasis, yet the high-resolution 3D remodeling of these networks under stress conditions remains poorly defined.

We employed Focused Ion Beam Scanning Electron Microscopy (FIB-SEM) to perform 3D reconstructions of INS1E cells subjected to mitochondrial respiratory chain inhibition, uncoupling, and exogenous oxidative stress. Quantitative analysis revealed that mitochondrial dysfunction induces profound ultrastructural transitions, characterized by significant luminal swelling of the ER, expansion of the perinuclear space, and mitochondrial diameter enlargement. 3D volume imaging identified a coordinated fragmentation of both ER and mitochondrial networks into discrete, spatially separated structures—a phenomenon distinct from the reticular morphology observed in control cells. The similarity between respiratory inhibition- and H_2_O_2_-induced phenotypes, together with preservation of ER structure following mitochondrial uncoupling, suggests a potential contribution of reactive oxygen species to the observed remodeling process.

Despite this extensive organelle breakdown, interorganelle membrane contact sites were not only preserved but expanded under stress conditions. We further provide a quantitative description of nuclear envelope–mitochondria contact sites (NAMs), demonstrating their selective remodeling during mitochondrial dysfunction. Our findings provide a high-resolution structural framework for organelle remodeling in β-cells, demonstrating that membrane contact sites are actively preserved and reorganized despite profound organelle fragmentation.

**Graphical abstract:** 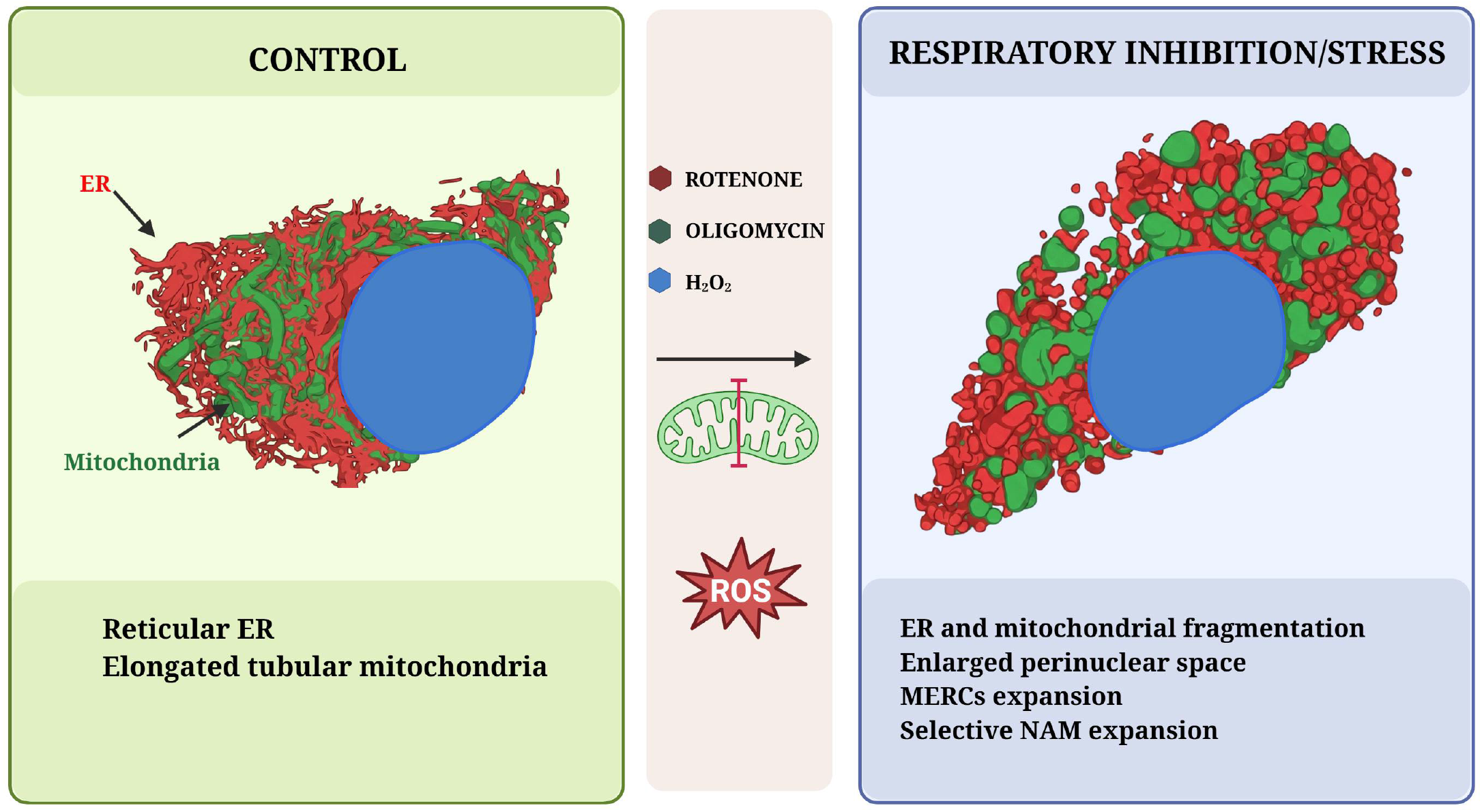

## Introduction

The maintenance of glucose homeostasis is critically dependent on the bioenergetic integrity of pancreatic β-cells, in which the mitochondrial respiratory chain serves as a central metabolic sensor coupling glucose metabolism to insulin secretion (Ježek et al., 2022; Maechler et al., 2006; Rutter et al., 2023). A growing body of evidence indicates that disruption of respiratory chain function markedly impairs both β-cell function and survival (Fujimoto et al., 2007; Maechler et al., 2010). Under conditions of chronic metabolic stress, including prolonged exposure to elevated glucose, mitochondrial respiration becomes compromised, leading to reduced ATP generation and defective glucose-stimulated insulin secretion (GSIS) (Chareyron et al., 2020; Ježek et al., 2012; Kabra et al., 2021). Dysfunction of specific respiratory chain components has been shown to be sufficient to impair insulin secretion and promote hyperglycemia in vivo (Aharon-Hananel et al., 2022; Lang et al., 2023). Beyond its role in energy production, impaired electron transport enhances the generation of reactive oxygen species (ROS), thereby disturbing redox homeostasis in β-cells, which possess relatively limited antioxidant capacity. Sustained oxidative stress activates apoptotic pathways and contributes to the progressive loss of functional β-cell mass (Bhatti et al., 2022; Dinić et al., 2022; Gerber and Rutter, 2017). Together, these findings establish respiratory chain integrity as a key determinant of β-cell performance and survival.

Disturbance of respiratory chain activity is closely associated with remodeling of mitochondrial morphology and dynamics (Bernard et al., 2007; Dlasková et al., 2010; Sauvanet et al., 2010). The mitochondrial network is highly dynamic and continuously undergoes fusion and fission to maintain functional integrity and adapt to changing metabolic demands (Chan, 2012; Ježek and Dlasková, 2019; Kawano et al., 2023; Rafelski, 2013). Mitochondrial fusion is mediated by mitofusin 1 and 2 (MFN1 and MFN2) at the outer mitochondrial membrane and optic atrophy 1 (OPA1) at the inner membrane, whereas fission is primarily driven by the recruitment of dynamin-related protein 1 (DRP1) to mitochondria via adaptor proteins such as FIS1 (Chen et al., 2003; Fonseca et al., 2019; Ishihara et al., 2006). The balance between these opposing processes is essential for maintaining mitochondrial network organization, cristae structure, and respiratory efficiency. Under chronic metabolic stress, this balance shifts toward increased fission and mitochondrial fragmentation, a phenotype associated with impaired oxidative phosphorylation and elevated ROS production (Adebayo et al., 2021; Chan, 2012; De Vos et al., 2005; Dlasková et al., 2010). Alterations in mitochondrial ultrastructure, including cristae remodeling, further compromise respiratory chain function and can facilitate the release of pro-apoptotic factors, thereby linking structural changes to β-cell dysfunction and cell death.

Compared to mitochondrial remodeling under stress conditions, relatively little is known about morphological changes of the endoplasmic reticulum (ER). In pancreatic β-cells, which have a high demand for insulin biosynthesis, disruption of ER homeostasis has profound functional consequences. Chronic ER stress activates the unfolded protein response (UPR), leading to translational attenuation and impaired proinsulin processing, resulting in an increased proinsulin-to-insulin ratio and defective insulin maturation (Diane et al., 2024; Kataoka and Noguchi, 2013; Makam et al., 2022). Morphologically, ER stress has been associated with cisternal dilation or ER “swelling,” reflecting altered membrane organization under conditions of secretory overload (Momose et al., 2006; Quan et al., 2012). Previous studies in Drosophila demonstrated that the ER possesses dedicated fusion and fission machinery that regulates its dynamic morphology (Espadas et al., 2019). Atlastin proteins mediate ER membrane fusion, whereas reticulons facilitate ER fission by constricting ER tubules. However, despite increasing knowledge of ER-shaping proteins, a detailed three-dimensional characterization of ER remodeling under stress conditions is still lacking.

Mitochondrial and endoplasmic reticulum (ER) stress are tightly coupled through coordinated signaling pathways, as well as through extensive physical and functional interactions between these organelles at mitochondria-associated ER membranes (MERCs) (Gottschalk et al., 2022; Kumar and Maity, 2021; van Vliet and Agostinis, 2018). These contact sites enable coordinated regulation of calcium signaling, lipid exchange, and cellular stress responses, allowing mitochondrial dysfunction to be transmitted to the ER (Bononi et al., 2012; Hajnóczky et al., 2000; Rusiñol et al., 1994; Sassano et al., 2022). A specialized domain of the ER is the nuclear envelope (NE), which surrounds the nucleus and also maintains direct contacts with mitochondria (Boulton-McDonald et al., 2026; Desai et al., 2020; Eisenberg-Bord et al., 2021). These nuclear envelope–mitochondria contacts (NAMs) are thought to contribute to retrograde signaling from mitochondria to nucleus, phospholipid metabolism and nuclear– cytoplasmic communication, although their regulation and functional significance remain poorly understood (Desai et al., 2020; Eisenberg-Bord et al., 2021).

Here, we demonstrate that remodeling of the mitochondrial network, driven specifically by respiratory chain dysfunction, is accompanied by a reorganization of the entire ER architecture. Furthermore, we show that respiratory chain perturbation not only remodels mitochondria–ER contact sites but also induces significant and previously uncharacterized changes in mitochondria–nuclear envelope contacts. These findings reveal an additional layer of cellular organization in which mitochondrial respiratory activity shapes the structural and functional landscape of ER-associated membranes. Collectively, our work positions mitochondrial bioenergetics and fitness as a central regulator of organelle crosstalk, providing a unifying mechanism linking metabolic stress to β-cell dysfunction and the progressive failure of insulin secretion.

## Methods

### Chemicals

All chemicals were obtained from Sigma-Aldrich unless stated otherwise.

### Cell culture

Rat insulinoma INS1E cells (AddexBio, San Diego, CA; cat. no. C0018009; RRID: CVCL_0351) were cultured as previously described (Kahancová et al., 2020). Cells were maintained in RPMI 1640 medium containing 11 mM glucose, 1 mM pyruvate, 2 mM L-glutamine, and 10 mM HEPES, supplemented with 5% (v/v) fetal calf serum, 50 μM β-mercaptoethanol, 50 IU/ml penicillin, and 50 μg/ml streptomycin. Cultures were kept at 37 °C in a humidified atmosphere with 5% CO_2_. Where indicated, cells were treated in standard culture medium with rotenone (5 µM, 2 h), oligomycin (1.25 µM, 2 h), H_2_O_2_ (250 µM, 2 h), FCCP (5 µM, 15 min), or left untreated (CTRL). These concentrations and treatment durations were selected based on conditions previously validated to induce respiratory chain inhibition or mitochondrial uncoupling.

### FIB-SEM microscopy

Samples were fixed for 2 h in a solution containing 2.5% glutaraldehyde and 2% paraformaldehyde prepared in 0.1 M cacodylate buffer (pH 7.2), followed by washing and postfixation in 2% OsO_4_ and 1.5% potassium ferrocyanide in the same buffer. An additional postfixation step was performed using 1% uranyl acetate in distilled water. Samples were then dehydrated through a graded ethanol series and embedded in EMBed-812 resin. After polymerization, blocks were trimmed, and regions of interest were mounted onto SEM stubs using conductive carbon and coated with a 25 nm platinum layer (Leica ACE600, High Vacuum Coater).

Imaging of INS1E cells was carried out using a Thermo Scientific Helios Hydra 5 CX system equipped with Auto Slice & View v. 5.7. Acquisition was controlled by FIB-SEM Maestro v. 0.2 (https://github.com/cemcof/FIBSEM_Maestro). Resin blocks were mounted on SEM stubs and sputter-coated with a 5 nm platinum layer (Quorum Q150T). A protective organoplatinum layer (~1.5 µm) was deposited over the region of interest using a gas injection system (30 kV, 13 pA). Trenches were milled with an oxygen plasma beam at 30 kV/5.6 nA, followed by polishing at 30 kV/0.61 nA and subsequent serial milling under identical conditions. Imaging was performed at 2 kV and 0.2–0.8 nA using an in-column backscattered electron detector (ICD) in immersion mode, with a voxel size of 5 or 10 nm and a dwell time of 5–15 µs.

### Data analysis

Images were binned to achieve an isotropic voxel size of 10 nm and converted to 8-bit depth using maximal histogram stretching. Subsequently, the dataset was aligned using the SIFT algorithm (Lowe, 2004) implemented in Fiji (Schindelin et al., 2012). Three-dimensional segmentation of mitochondria, endoplasmic reticulum (ER), nuclei, and membrane contact sites was performed in either Microscopy Image Browser (MIB) (Belevich et al., 2016) or NIS-Elements using the AI segmentation module, followed by manual inspection and correction of segmented volumes. ER segmentation captured predominantly well-resolved cisternal and tubular regions. Thin ER tubules approaching the resolution limit of the dataset were not consistently traceable throughout the reconstructed volume and were therefore only partially segmented. Consequently, the reconstructed ER network represents a partial segmentation of confidently identifiable ER structures.

Surface renderings and volumetric measurements were generated using marching cubes-based surface reconstruction. Quantitative analyses were restricted to fully resolved structures contained within the analyzed volume to minimize edge-related bias. ER–mitochondria contact sites (MERCs) and nucleus-associated mitochondrial contact sites (NAMs) were defined as membrane regions separated by ≤30 nm. Surface area, object volume, and contact site area measurements were extracted from segmented 3D objects. Equivalent sphere diameter values were calculated from segmented object volumes to estimate the size distribution of fragmented ER structures.

For mitochondria-associated contact site analysis, individual mitochondrial objects were identified by three-dimensional connected-component labeling using 26-connectivity. Mitochondrial surface area was estimated from binary mitochondrial masks using marching cubes surface reconstruction with isotropic voxel spacing of 10 × 10 × 10 nm. Contact site masks were intersected with mitochondrial surfaces to restrict measurements to contact regions localized at the mitochondrial boundary. Individual contact domains were subsequently separated by connected-component analysis and quantified per mitochondrial object, including contact surface area and the percentage of mitochondrial surface occupied by contact sites.

Individual NAM contact regions were extracted from binary contact masks and analyzed as three-dimensional connected components. Connected-component labeling was performed using 26-connectivity. For each connected NAM component, surface area was quantified. Surface area was calculated from binary component masks using marching cubes reconstruction, followed by mesh-based surface area extraction.

Surface renderings for three-dimensional visualization were generated in Amira software using marching cubes-based surface reconstruction.

### Statistical analysis

Statistical analyses and graphical visualization were performed using GraphPad Prism software. Quantitative data are presented as mean ± S.D. unless otherwise stated. Outliers were identified using the 1.5 × interquartile range (IQR) criterion and excluded prior to statistical evaluation. Statistical significance was assessed using the tests indicated in the corresponding figure legends.

## Results

### Inhibition of mitochondrial respiration remodels ER and nuclear envelope architecture in INS1E cells

To examine the impact of respiratory chain inhibition on the 3D architecture of mitochondria, the endoplasmic reticulum (ER), and their contact sites in the model pancreatic β-cell line INS1E, we employed focused ion beam scanning electron microscopy (FIB-SEM). Cells were treated with rotenone (a complex I inhibitor), oligomycin (an ATP synthase inhibitor), or FCCP (a mitochondrial uncoupler), alongside H_2_O_2_ as a non-mitochondrial oxidative stress control (Fig. 1). Quantitative analysis of 2D sections demonstrated distinct, treatment-specific alterations in ER morphology (Fig. 1B). Rotenone, H_2_O_2_, and oligomycin induced profound ER luminal swelling, increasing the average ER diameter from 46 nm in control cells to 339 nm, 471 nm, and 291 nm, respectively. Conversely, FCCP treatment increased ER diameter only to 58 nm, without markedly changing the ER 2D morphology.

**Figure 1.**
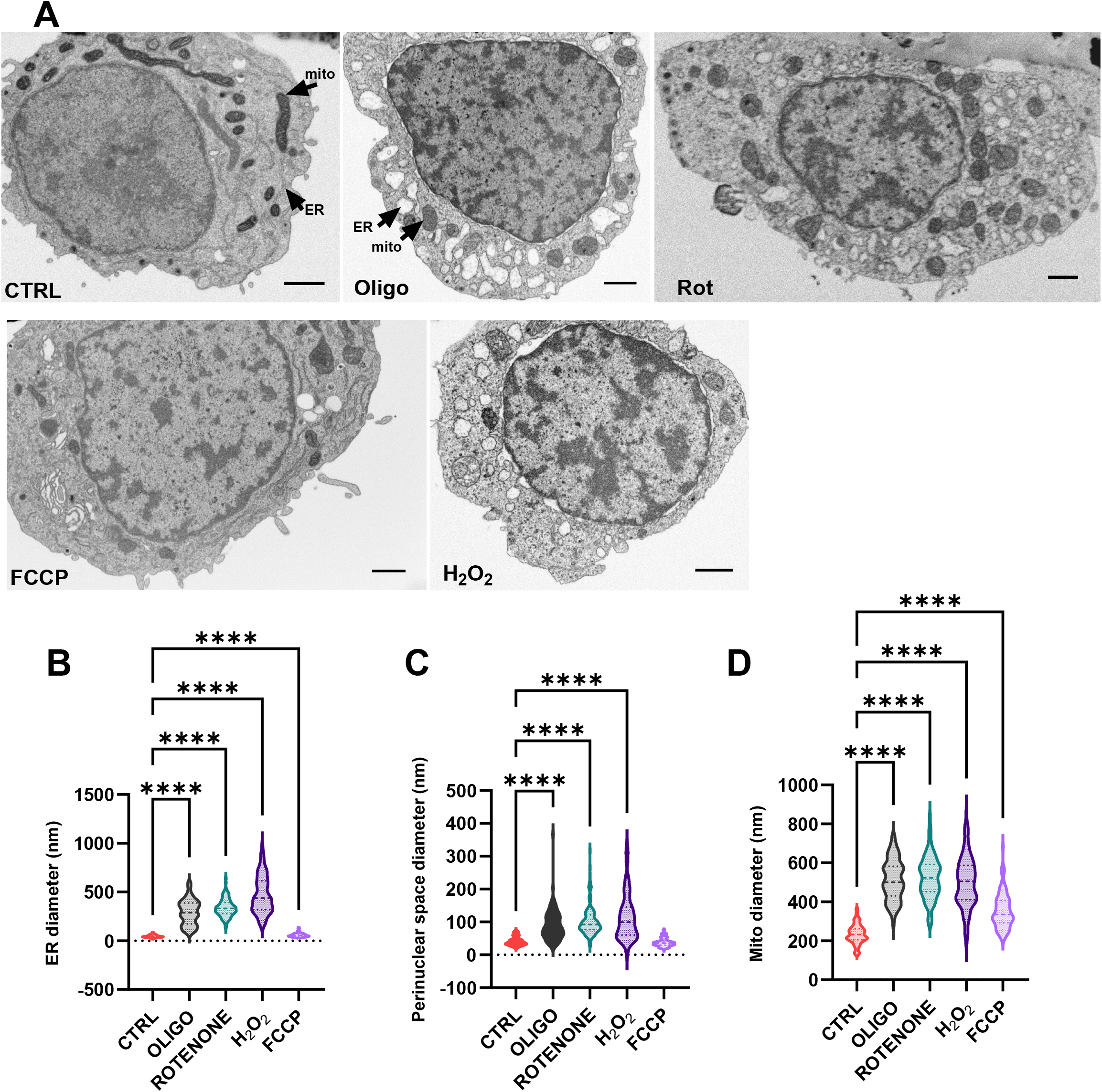
2D ultrastructural analysis reveals mitochondrial stress-induced remodeling of ER, perinuclear space, and mitochondria in INS1E Cells. (A) Representative FIB-SEM micrographs of INS1E cells treated with rotenone (5 µM, 2 h), oligomycin (1.25 µM, 2 h), H_2_O_2_ (250 µM, 2 h), FCCP (5 µM, 15 min), or left untreated (CTRL). Mitochondrial stress induction resulted in pronounced ultrastructural alterations of intracellular membranes and mitochondrial morphology. *Mito*, mitochondria; *ER*, endoplasmic reticulum. Scale bar: 1000 nm. (B) Quantification of ER tubule/fragment diameter. (C) Quantification of perinuclear space diameter. (D) Quantification of mitochondrial diameter. Violin plots represent the distribution of measured values for each treatment condition (n = 100 analyzed diameters per condition). Statistical significance was evaluated using Welch’s ANOVA with the Games-Howell post hoc test. ****P < 0.0001. Data are presented as means ± S.D.

Parallel changes were observed in the perinuclear space (Fig. 1C). Rotenone, oligomycin, and H_2_O_2_ increased the diameter of the perinuclear space from approximately 41 nm in control cells to 104 nm, 90 nm, and 114 nm, respectively. In contrast, FCCP treatment did not significantly change the diameter.

Finally, mitochondrial morphology closely mirrored these ultrastructural trends (Fig. 1D). Rotenone, oligomycin, and H_2_O_2_ increased mitochondrial diameter from approximately 235 nm in control cells to 524 nm, 513 nm, and 503 nm, respectively. FCCP treatment also increased mitochondrial diameter to 356 nm, although to a lesser extent than the other stress conditions.

Taken together, these data demonstrate extensive, coordinated remodeling of the ER, nuclear envelope, and mitochondrial architecture in response to mitochondrial stress. Furthermore, the phenotypic similarity between the rotenone-, oligomycin-, and H_2_O_2_-treated groups suggests that oxidative stress might be a key driver of this structural remodeling.

### 3D FIB-SEM reveals coordinated fragmentation of ER and mitochondrial networks

Three-dimensional FIB-SEM reconstruction demonstrated extensive remodeling of both the ER and mitochondrial networks following mitochondrial respiratory inhibition (Fig. 2). Control cells displayed a thin reticular ER network closely associated with elongated tubular mitochondria. In contrast, both inhibition of the mitochondrial respiratory chain and external oxidative stress induced by H_2_O_2_ led to fragmentation of both organelle systems, characterized by conversion of mitochondria and ER into rounded, fragmented structures. Quantitative analysis of fragmented ER components revealed equivalent sphere diameters ranging from approximately 158–218 nm across treatment conditions, with no significant differences observed between oligomycin-, rotenone-, and H_2_O_2_-treated cells (Fig. 3). In contrast, mitochondrial uncoupling by FCCP largely preserved the ER network, while mitochondria retained a more elongated morphology with thickened tubules and occasional donut-like structures.

**Figure 2.**
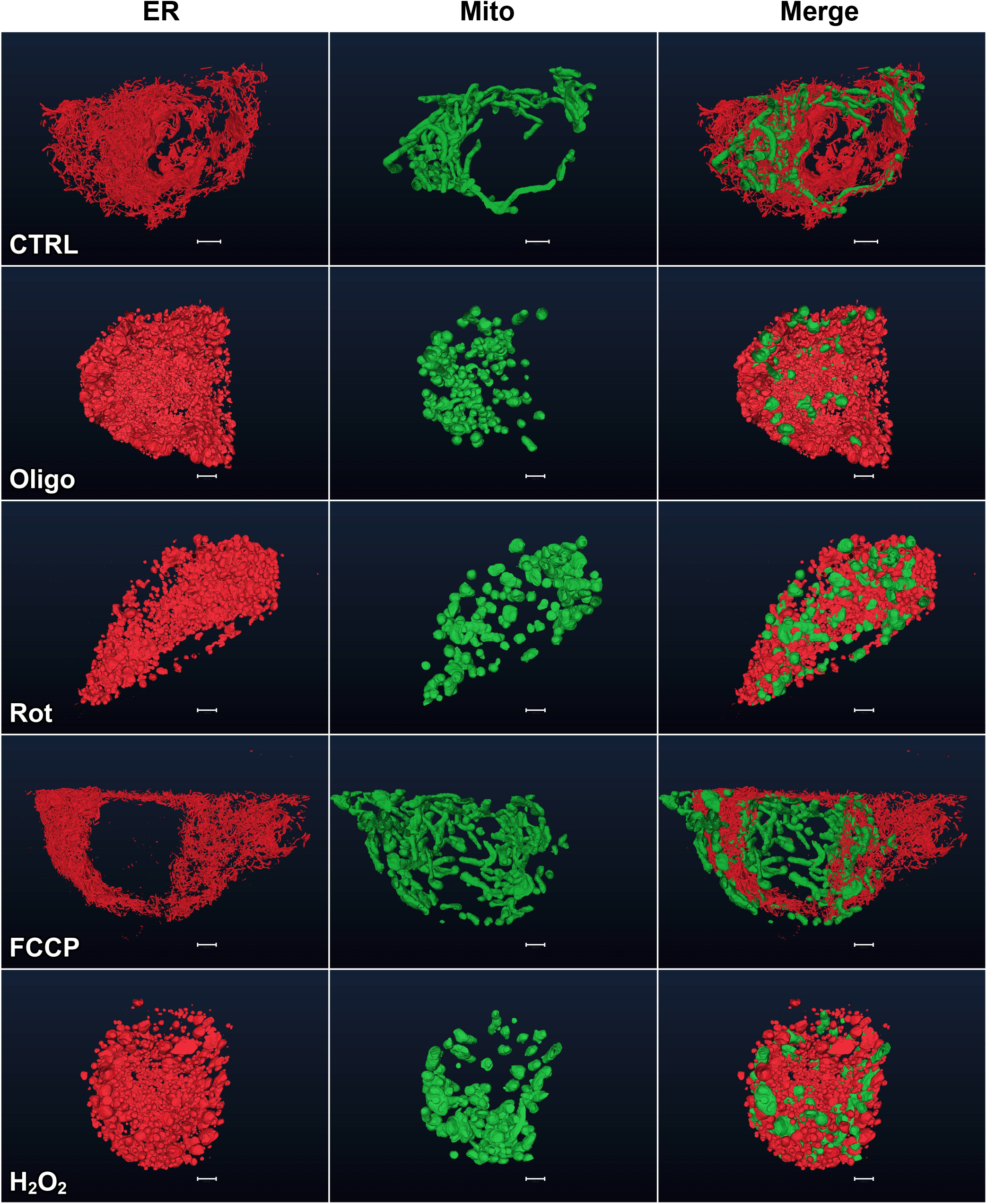
3D reconstruction of ER and mitochondrial networks reveals stress-induced organelle remodeling in INS1E cells. Representative 3D surface renderings of segmented endoplasmic reticulum (ER, red) and mitochondria (mito, green) in INS1E cells left untreated (CTRL) or treated with oligomycin (1.25 µM, 2 h), rotenone (5 µM, 2 h), FCCP (5 µM, 15 min), or H_2_O_2_ (250 µM, 2 h). Rows correspond to individual treatment conditions, while columns show ER segmentation, mitochondrial segmentation, and merged ER–mitochondria reconstructions. The merged images illustrate the spatial organization of mitochondria relative to the ER network under control and mitochondrial stress conditions. Scale bars are 1000 nm.

**Figure 3.**
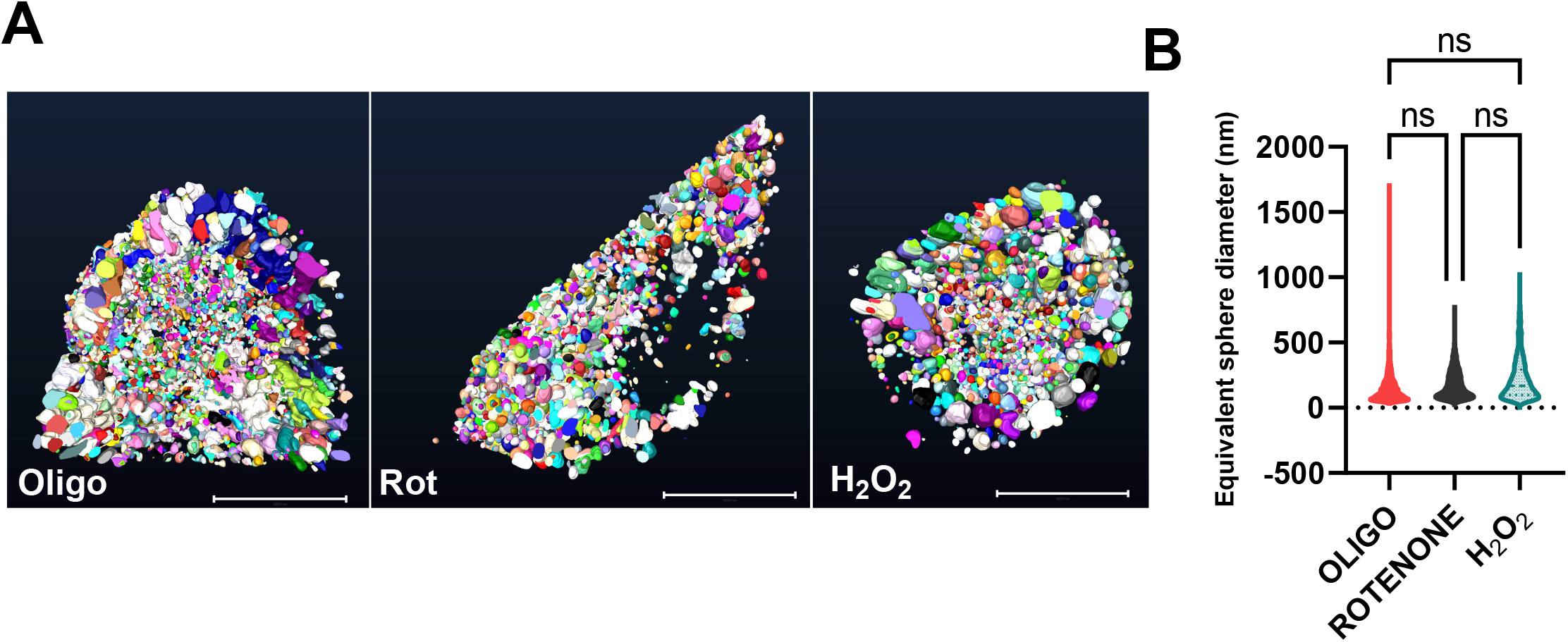
Analysis of fragmented ER structures in INS1E cells. (A) Representative 3D surface renderings of segmented endoplasmic reticulum (ER) structural components in INS1E cells treated with oligomycin (1.25 µM, 2 h), rotenone (5 µM, 2 h), or H_2_O_2_ (250 µM, 2 h). Different colors represent individual segmented ER components within the reconstructed ER network, illustrating treatment-dependent alterations in ER organization and morphology. Scale bars are 5000 nm. (B) Quantification of ER component size represented as equivalent sphere diameter (nm) for each treatment condition. Violin plots show the distribution of ER component diameters across analyzed cells (n > 1000 analyzed ER fragments per condition). Statistical significance was evaluated using Welch’s ANOVA with Games-Howell post hoc test. No significant differences (ns) were observed between treatment groups. Data are presented as means ± S.D.

Together, these data characterize dramatic remodeling of 3D ER and mitochondrial architecture in response to mitochondrial dysfunction and oxidative stress.

### ER–mitochondria contact sites are preserved despite organelle fragmentation

Despite the extensive fragmentation of both ER and mitochondrial networks, mitochondria–ER contact sites (MERCs) remained preserved across all treatment conditions (Fig. 4). Three-dimensional FIB-SEM reconstructions revealed persistent ER–mitochondria appositions in control, oligomycin-, rotenone-, FCCP-, and H_2_O_2_-treated cells. Although both organelles adopted fragmented morphologies following respiratory inhibition and oxidative stress, close membrane contacts between the ER and mitochondria were still frequently observed.

**Figure 4.**
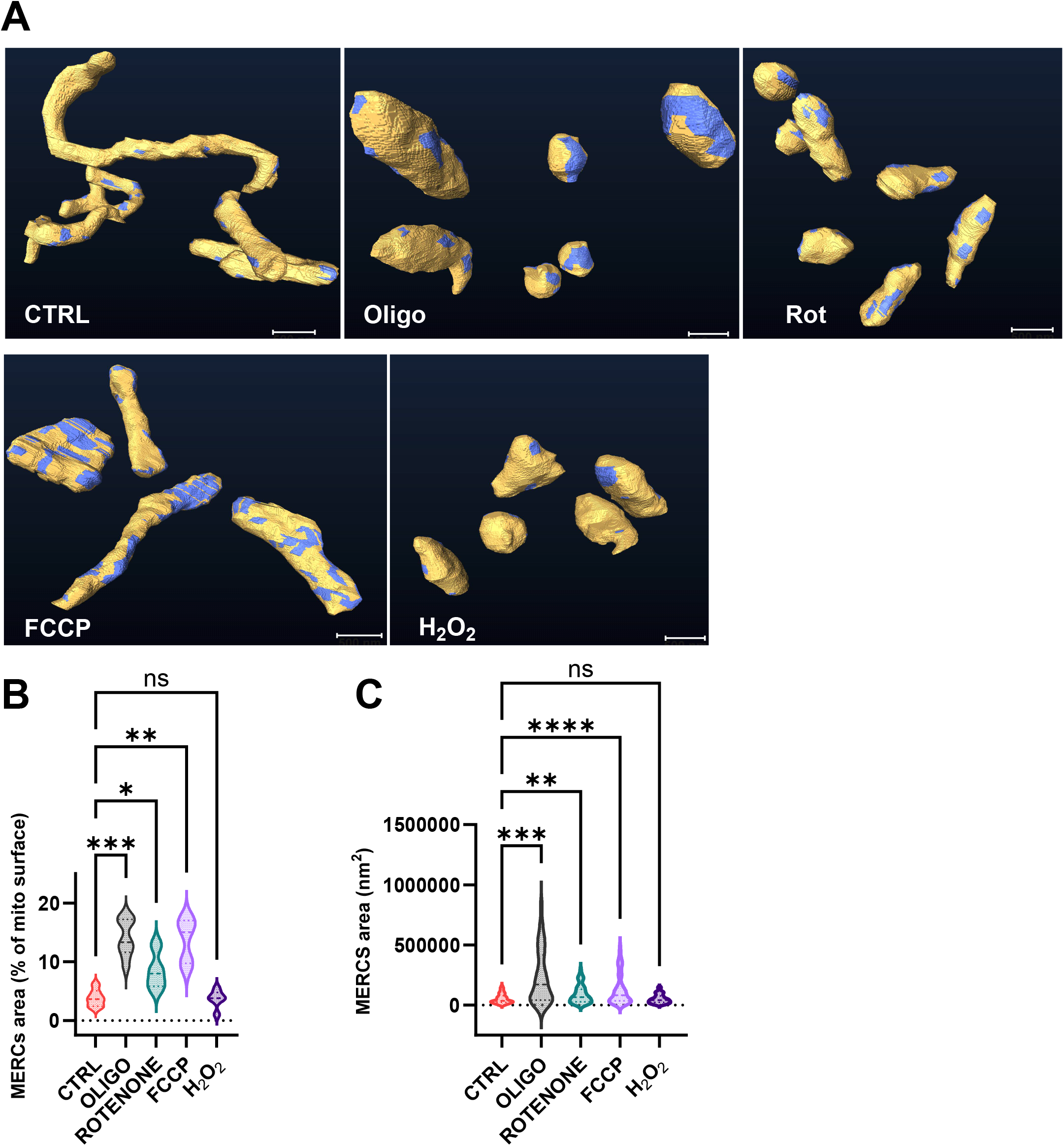
Mitochondrial stress alters ER–mitochondria contact site abundance and area in INS1E cells. (A) Representative 3D surface renderings of segmented mitochondria in INS1E cells treated with rotenone (5 µM, 2 h), oligomycin (1.25 µM, 2 h), H_2_O_2_ (250 µM, 2 h), FCCP (5 µM, 15 min), or left untreated (CTRL). Mitochondria are shown in yellow, and ER–mitochondria contact regions are highlighted in blue. Contact sites were defined as ER membrane regions located within 30 nm of the mitochondrial surface. Scale bars: 500 nm. (B) Quantification of ER–mitochondria contact site abundance, expressed as the percentage of MERC area relative to total mitochondrial surface area (n > 5 analyzed mitochondria per condition). (C) Quantification of total MERC area in nm^2^ (n > 30 analyzed MERCs per condition). Violin plots represent the distribution of measured values for each treatment condition. Statistical significance was evaluated using Welch’s ANOVA with the Games-Howell post hoc test. Statistical significance is indicated in the graphs: *P < 0.05, **P < 0.01, ***P < 0.001, ****P < 0.0001; ns, not significant. Data are presented as means ± S.D.

Quantitative analysis demonstrated that the proportion of mitochondrial surface associated with MERCs increased from approximately 4% in control cells to 14% following oligomycin treatment, 8% following rotenone treatment, and 14% following FCCP treatment (Fig. 4B). In contrast, H_2_O_2_ treatment did not alter MERC abundance, remaining at approximately 4% despite substantial ER and mitochondrial fragmentation. Analysis of absolute MERC surface area revealed a similar trend, with mitochondrial stress conditions leading to a marked expansion of ER–mitochondria contact interfaces, particularly following oligomycin and FCCP treatment, whereas H_2_O_2_ induced only minimal changes.

Notably, ER–mitochondria contacts remained preserved despite extensive fragmentation of both organelle systems and were partially expanded under conditions of mitochondrial dysfunction.

### Nuclear envelope–mitochondria contact sites are remodeled under mitochondrial stress conditions

Three-dimensional FIB-SEM reconstruction revealed that nuclear envelope–mitochondria contact sites (NAMs) undergo substantial remodeling in response to mitochondrial stress (Fig. 5). In control cells, NAMs appeared as discrete and localized contact regions between mitochondria and the nuclear envelope, occupying approximately 4% of the nuclear surface area. However, mitochondrial inhibition and oxidative stress altered the extent of these associations. Quantitative analysis of NAM surface area demonstrated an overall trend toward expansion following stress, although the magnitude of this effect was treatment-dependent (Fig. 5). Oligomycin and H_2_O_2_ induced the most pronounced increases in NAM area compared to control cells (P < 0.0001 and P < 0.05, respectively). In contrast, rotenone and FCCP produced only minor shifts in NAM area distribution that did not reach statistical significance. Despite extensive remodeling of mitochondrial and ER networks, close nuclear envelope–mitochondria contacts associations remained preserved across all treatment conditions.

**Figure 5.**
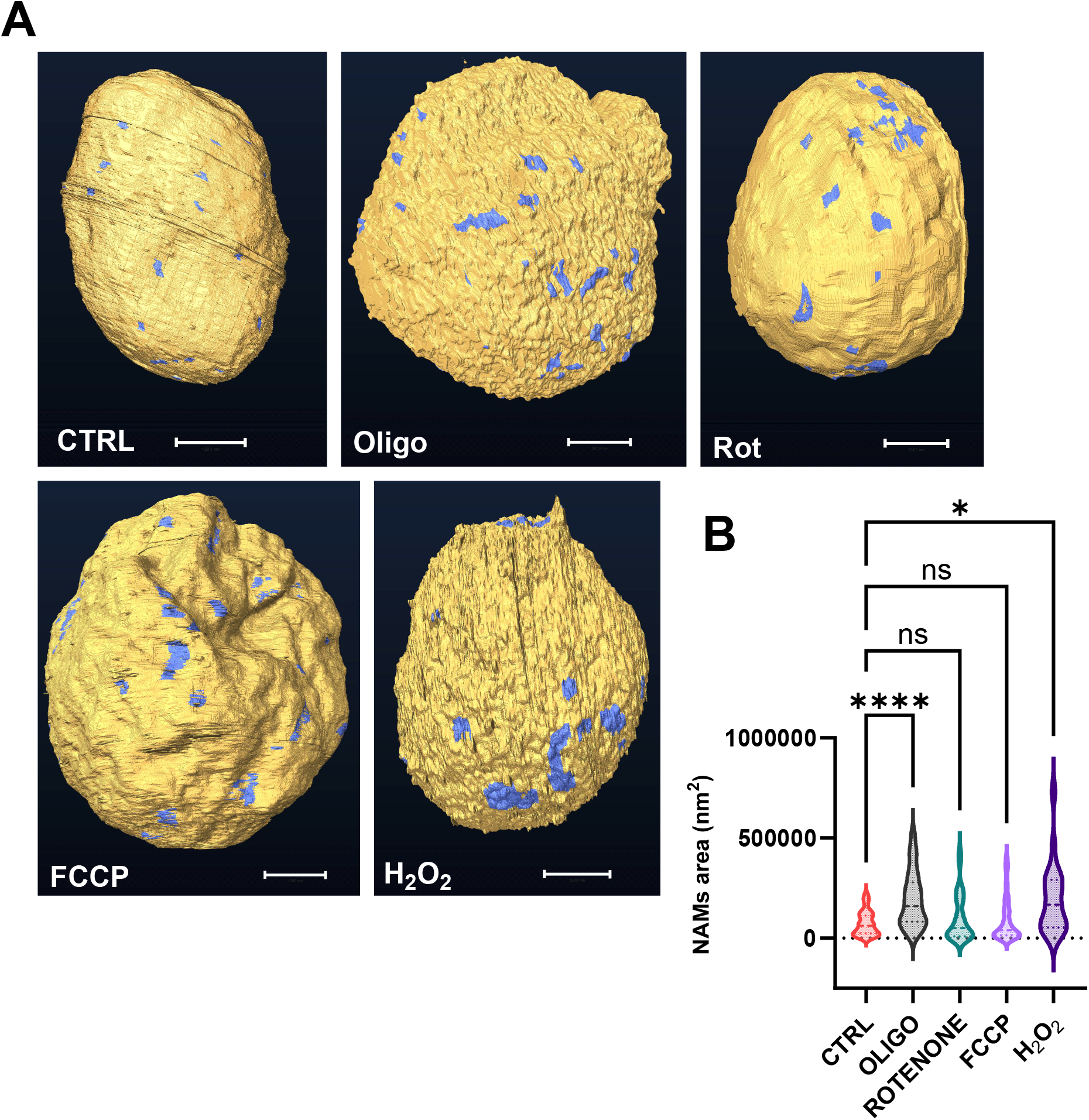
Mitochondrial stress alters nucleus-associated mitochondrial contact areas in INS1E cells. (A) Representative 3D surface renderings of segmented nuclei from INS1E cells treated with rotenone (5 µM, 2 h), oligomycin (1.25 µM, 2 h), H_2_O_2_ (250 µM, 2 h), FCCP (5 µM, 15 min), or left untreated (CTRL). Nuclear surfaces are shown in yellow, and nucleus-associated mitochondrial contact regions are highlighted in blue. Contact sites were defined as nuclear membrane regions located within 30 nm of the mitochondrial surface. Scale bars: 1500 nm. (B) Quantification of nucleus-associated mitochondrial contact site area, expressed as NAM area in nm^2^ (n > 20 analyzed NAMs per condition). Violin plots represent the distribution of measured values for each treatment condition. Statistical significance was evaluated using Welch’s ANOVA with Games-Howell post hoc test. Statistical significance is indicated in the graph: *P < 0.05, ****P < 0.0001; ns, not significant. Data are presented as means ± S.D.

Collectively, these data indicate that the NAM interface is a dynamic structural feature that selectively expands under specific forms of mitochondrial dysfunction, particularly during ATP synthase inhibition and exogenous oxidative stress.

## Discussion

In pancreatic β-cells, the endoplasmic reticulum (ER) is exceptionally sensitive to cellular stress due to its high secretory load and central role in insulin synthesis. To preserve functional integrity and support β-cell survival, the ER deploys multiple adaptive mechanisms that maintain protein folding capacity, calcium homeostasis, lipid synthesis and membrane organization. Under stress conditions, β-cells undergo profound ultrastructural remodeling that reflects both adaptive and maladaptive responses. In conventional 2D electron micrographs, this stress phenotype is characterized primarily by swollen ER cisternae, an enlarged perinuclear space, and enlarged mitochondria. However, the spatial organization of these structural changes within the three-dimensional cellular context has remained insufficiently understood.

Using 3D volume electron microscopy, our study provides a detailed ultrastructural characterization of ER reorganization induced by respiratory chain inhibition and oxidative stress. We demonstrate that mitochondrial dysfunction is accompanied by coordinated fragmentation of both mitochondrial and ER networks, resulting in the breakdown of the normally thin ER network into disconnected and spatially separated structures. Although ER fragmentation has previously been suggested in neurons and other cell types using fluorescence microscopy, those observations were largely inferred indirectly from discontinuities in fluorescence signals or cytosolic voids (Bao et al., 2016; Kucharz et al., 2013, 2011, 2009). The volumetric ultrastructural analysis presented here directly reveals the extent and spatial organization of ER fragmentation within intact β-cells, demonstrating that ER remodeling under stress is substantially more extensive than can be appreciated from conventional 2D imaging alone. Whether this pronounced ER fragmentation reflects a general stress response or represents a specialized feature linked to the highly active secretory nature of pancreatic β-cells remains an intriguing question for future studies.

The observed remodeling appears to depend strongly on the precise nature of the mitochondrial stressor. Inhibition of respiration by rotenone or oligomycin, as well as exogenous oxidative stress induced by H_2_O_2_, produced remarkably similar phenotypes characterized by ER swelling, loss of ER architecture, mitochondrial fragmentation, and the expansion of membrane contact sites. In contrast, mitochondrial uncoupling by FCCP largely preserved ER network integrity and sustained less fragmented mitochondrial morphology (although a small subset adopted a donut-like morphology), suggesting that proton motive force dissipation alone distinctively alters organelle dynamics compared to respiratory arrest. The strong phenotypic similarity between respiratory inhibition and H_2_O_2_ treatment suggests that reactive oxygen species (ROS) might contribute substantially to the observed ER remodeling.

An important finding of our study is that the extensive organelle fragmentation did not disrupt membrane contact site formation. Mitochondria–endoplasmic reticulum contact sites (MERCs) remained preserved under all treatment conditions and were even expanded following specific forms of mitochondrial stress. These observations support the emerging paradigm that membrane contact sites are highly robust, dynamically regulated structures that persist independently of global organelle morphology and fusion/fission machinery. Increased MERC abundance under stress conditions may reflect an adaptive cellular response aimed at maintaining residual calcium transfer, optimizing lipid exchange, or facilitating interorganelle stress signaling. Previous studies have demonstrated that ER stress and mitochondrial dysfunction can promote the assembly of molecular tethering complexes at MERCs (Casas-Martinez et al., 2025; Shiiba et al., 2025); our data extend these observations by visually demonstrating that MERCs persist even when both mitochondrial and ER networks undergo severe, macroscale fragmentation.

Beyond MERCs, our study quantitatively characterizes nucleus-associated mitochondrial contact sites (NAMs), a structurally and functionally elusive category of membrane contact sites that has remained technically challenging to analyze in three dimensions. Although previous reports described the pooling of mitochondria in the perinuclear region during stress (Agarwal and Ganesh, 2020; Al-Mehdi et al., 2012), such clustering does not inherently imply the formation of direct, functional membrane interfaces. By combining FIB-SEM reconstruction with quantitative surface analysis, we demonstrate that NAM interfaces selectively expand during specific forms of mitochondrial dysfunction, particularly following ATP synthase inhibition and oxidative stress. These observations establish a quantitative ultrastructural framework for studying NAM remodeling under metabolic stress, potentially pointing toward an augmented retrograde signaling axis or localized ATP provisioning to the nucleus. The preservation and expansion of NAMs despite extensive organelle fragmentation further emphasize that membrane contact site remodeling is an active, coordinated program rather than a passive consequence of stochastic organelle apposition.

In conclusion, our study provides a comprehensive three-dimensional ultrastructural analysis of mitochondrial stress-induced remodeling in pancreatic β-cells. We identify ER fragmentation as a prominent structural hallmark of respiratory chain inhibition and oxidative stress, and demonstrate that membrane contact sites remain preserved and dynamically reorganized despite severe disruption of global organelle architecture. These findings expand our current understanding of organelle plasticity during mitochondrial dysfunction and provide a framework for future studies investigating how disrupted interorganelle communication drives β-cell failure in diabetes.

## CRediT authorship contribution statement

Andrea Dlasková: Conceptualization, Investigation, Data curation, Supervision, Funding acquisition, Writing – original draft. Bazila Bazila: Conceptualization, Investigation, Data curation. Pavel Křepelka: Investigation, Data curation. Rhea Christine Victor: Data processing. Darsh Jaresh Jhala: Data processing. Petr Ježek: Writing – review & editing..

## Acknowledgements

The authors acknowledge the Imaging Methods Core Facility at IMG, supported by the Ministry of Education, Youth and Sports of the Czech Republic (LM2023050 Czech-BioImaging), for technical support and assistance. We further acknowledge CF CryoEM of CIISB, Instruct-CZ Center, supported by MEYS CR (LM2023042) and European Regional Development Fund-Project “Innovation of Czech Infrastructure for Integrative Structural Biology” (No. CZ.02.01.01/00/23_015/0008175).

